# Aperiodicity in mouse CA1 and DG power spectra

**DOI:** 10.1101/2025.01.30.635678

**Authors:** Gustav Kühn, Hussin El Rashidy, Federico Calegari, Shonali Dhingra

## Abstract

Rodent hippocampal power spectra comprise of periodic and aperiodic components. The periodic components (brain rhythms) contain information about the behavioral or cognitive state of the animal. The aperiodic components are rarely studied and their functionality is not well understood, though have shown to be correlated with animal’s age or the excitation-inhibition ratio of the brain region. To study these components in the mouse hippocampus we modified the existing open-source FOOOF toolbox, which was originally optimized for EEG data. First, using simulated data, we show that our modifications decrease the error in assessment of the periodic components from 3% to 0.1%. Second, using tetrode electrophysiological signals, we compare the aperiodic activity within mice hippocampal sub-regions, CA1 and dentate gyrus (DG). Our optimization of FOOOF improved the aperiodic assessment errors by about 50% and were critical in making the first-ever assessments of the aperiodic components in these brain regions. Our results show that the aperiodic parameters in the DG are multi-fold and significantly larger than in CA1, revealing higher excitation and longer timescales in CA1 than DG. Our work highlights the subtle differences in electrophysiology field potentials between hippocampal sub-regions, and presents the improvements needed in the existing open-source toolbox to be able to see such differences.

## INTRODUCTION

EEG is one of the oldest and most widely used methods for the investigation of the electrical activity of the brain. On the other hand, *in-vivo* electrophysiology (ephys) provides wide-band neural signals (direct current to 40kHz), which contain both action potentials and other membrane potential-derived fluctuations in a small neuronal volume, termed as local field potential (LFP). The scalp EEG is a spatiotemporally smoothed version of the LFP, integrated over an area of 10cm^2^ or more^1^. Though LFP in ephys signals provides a more detailed view of the local neuronal environment, being more invasive than EEG makes it less commonly used in human studies, but frequently employed in rodent studies.

The wideband LFP signals contain oscillatory activity overlaid on top of aperiodic background. Neural oscillations are prominent features of rhythmic brain activity, which are thought to relate to population synchrony and coordination of neural activity at the population level2–5. The most prevalent of these brain rhythms in the rodent hippocampus include theta (4-12Hz) and slow gamma (sgamma, 25-50Hz)^6^. Along with these brain rhythms, the power distribution at various frequencies follows a 1/f^n^ statistics, commonly attributed to various extra-somatic and extracellular sources^1^. This aperiodic roll-off with frequency in these signals is occasionally cast aside as “noise”, due to its ubiquitous nature. Investigation of the aperiodic components has recently gained considerable interest, and the exponent *n* has been shown to change with task demands, age, and disease, among other correlations^1,4,7,8^. Though, to the best of our knowledge, complete estimation of the aperiodic components in the rodent hippocampus has never been reported.

Traditionally, the most commonly used method to estimate parameters of oscillatory activity in ephys data has been bandpass filtering^9^. Due to the above-mentioned aperiodic activity, applying narrowband filtering can lead to nonzero power in the detection bands^4,10^. This is sometimes avoided by broader band filtering and normalization by the average broad-band power^11^, which could still lead to biased estimates of power in a certain narrow band, and frequency of maximum power in such a band, especially for lower frequency bands (supp fig 1a).

Recently, there has been a big thrust in determining the aperiodic components of the neural signals to get better estimates of the periodic components^12^, and correlate them with behavioral or cognitive measures, such as mentioned above. A recently developed method, termed FOOOF, is currently widely used to separate out the rhythmic neural oscillations, from the aperiodic or arrhythmic part of the spectrum^13^. This analytical method has mostly been used for EEG and ECoG studies in humans and macaques, with only 2 reports so far using it to analyze hippocampal ephys recordings in rodents^14,15^.

To be able to reliably use FOOOF for hippocampal ephys signals, we made vital modifications to its assessment procedures of aperiodic and periodic components (see results and methods for details). In brief, we added 3 new aperiodic fitting models more suited for ephys signals, restricted the detection of the periodic components to be within hippocampus-relevant frequency bands, and iteratively implemented these steps to minimize errors. We then used our modified toolbox to analyze recordings in the CA1 and dentate gyrus (DG) of the mouse hippocampus.

## RESULTS

### Improved assessment of periodic parameters

Over the years, several methods have been used to estimate the parameters of various periodic components in ephys LFP. Most of these methods can be categorized into one of the two methodologies of either working in the time-domain or in the frequency-domain. For the methods working in the time-domain, the time series data is subjected to either narrow^16^ or broad^17^ bandpass filtering, further using the filtered data to estimate the center frequency and power in a specific frequency band. In the more recent papers, with the understanding of the existence of 1/f^n^ statistics in the neural data, these estimates were made on the data in the frequency domain^9,11,18^.

To test the efficacy of the commonly used time-domain and frequency-domain methods in estimating the center frequency of hippocampal theta rhythm, we simulated a Gaussian theta peak with a center frequency (cf) of 6-9Hz, and a standard deviation of 1-2.5Hz (2-5Hz bandwidth (bw)). To insert an aperiodic component into the same, we added a ‘pink noise-like’ spectrum into it, which is simply a 1/f^n^ noise, with an exponent of 1.2. For a realistic simulation, we also added a noise peak at 50Hz (see methods for more details, fig 1a). The parameters for simulations were chosen based on the medians in real tetrode ephys data from the mouse hippocampus.

**Fig 1:**
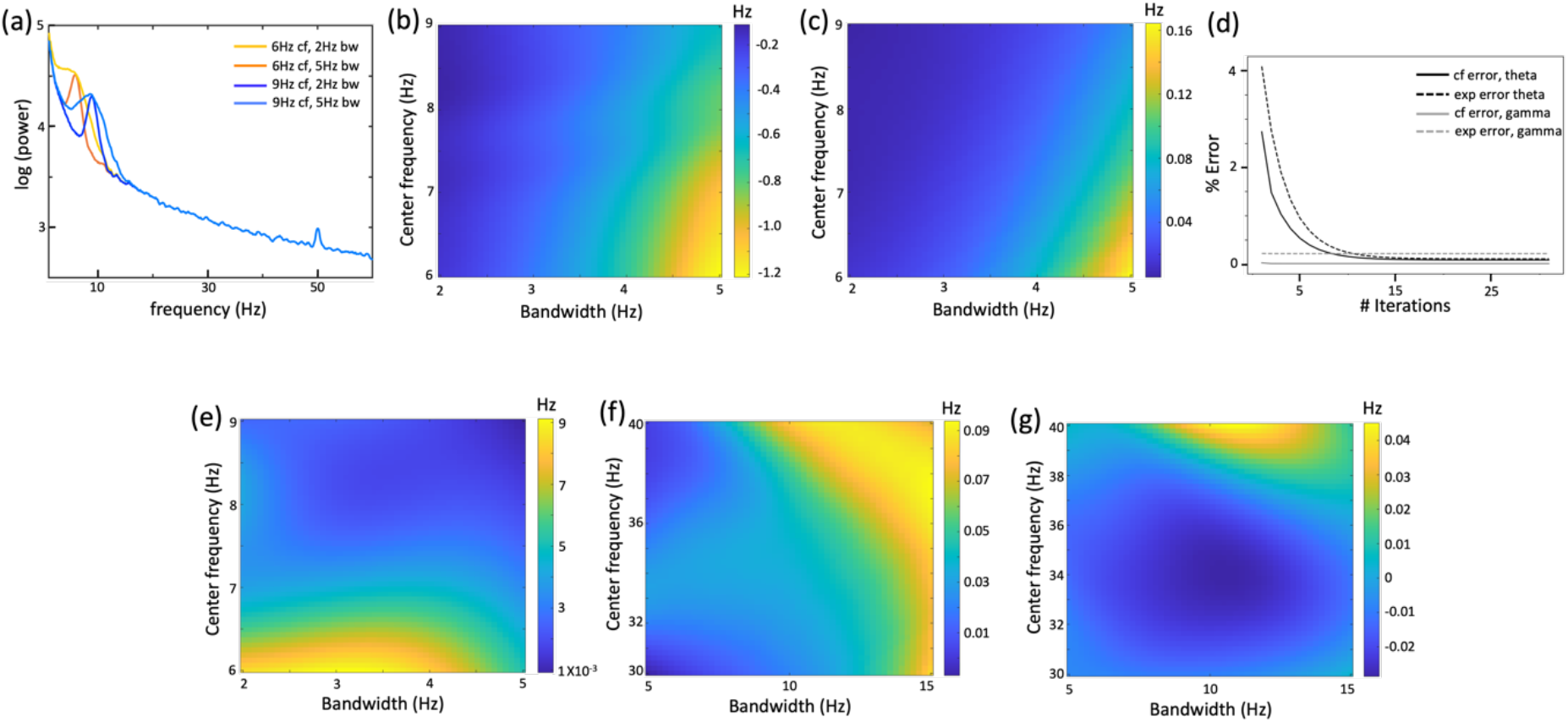
Better estimates of the periodic components in simulated electrophysiological signals. (a) 4 example simulated hippocampal signals, with theta as a Gaussian peak with a center frequency (cf) varying between 6-9Hz, bandwidth (bw) between 2-5Hz, and power of about 15dB, with pink-noise exponent of 1.2. Generated signals spanned 1-100Hz, but shown between 1-60Hz here for better visualization. (b) Errors in cf estimates of the simulated theta peak using Chronux toolbox, with varying bw and cf. This and further such plots were smoothed using a Gaussian filter with std of 4, unless specified. (c) Average errors in cf estimates using FOOOF, across 20 randomizations, of the simulated theta peak, with varying bw and cf. (d) Percentage errors in cf and exponent estimates for simulated theta (cf=6Hz, bw=5Hz) and sgamma (cf=30Hz, bw=15Hz) peaks with iterative aperiodic and periodic estimation in FOOOF. (e) Same as (c) after 20 iterations in FOOOF. (f,g) Same as (c,e) but for simulated sgamma peaks.

Using both narrow bandpass filtering (in time-domain) and chronux toolbox (in frequency-domain) led to 15-20% error in the cf estimates of the theta Gaussian peak (fig 1b, supp fig 1a). These estimates were comparably erroneous when estimated in the time-domain or frequency-domain, with the center frequencies mostly getting underestimated. Further, as expected, the errors in these estimates increase with bw and decreasing cf, and decrease for smaller exponents (fig 1b vs. supp fig 1b).

A reported advantage of using FOOOF toolbox is to get unbiased estimates of various periodic components, including maximum power, cf and bw. A suggested way to use FOOOF is to subtract the estimated aperiodic component, and use the ‘flattened’ spectrum to estimate the periodic components by fitting Gaussian peaks^14^. Using FOOOF this way on the simulated data, the errors in cf estimates indeed went down to about 3% (fig 1c), but the power and cf estimates were still visibly inaccurate (supp fig 1d).

To improve the periodic parameter estimates with FOOOF, we modified it to iteratively estimate the aperiodic components to obtain a flattened spectrum, fitting Gaussians to the periodic components, and subtracting the oscillatory parts from the original spectrum to estimate the aperiodic components again. The errors in estimates of the cf and the exponent of the aperiodic component decreased with the iterations (fig 1d), for the simulated theta peaks. After 20 iterations, the errors in the cf estimates dropped down to less than 0.1% (fig 1d,e). To simulate peaks in the sgamma band, we put in a Gaussian peak with cf of 30-40Hz, and bw of 5-15Hz. As expected, the iterative procedure does not affect the cf and exponent estimates of sgamma as much (fig 1d), and decreases the cf error estimates from 0.3% to 0.0001% (fig 1f,g).

### Improved aperiodic fitting until 200Hz

Beyond estimating the periodic components, we further used FOOOF to analyze the aperiodic components of ephys signals from the mouse hippocampus. FOOOF allows doing this by fitting a linear 1/f^n^ function to the signals (see methods, equation (1)), giving the exponent *n* as a possible signature of the acquisition system or biologically relevant parameters. The exponent *n* has usually been estimated within a small range of frequencies, with recent reports arguing that *n* would depend on the frequency range of estimation^19^. This exponent has been modeled in the range of 30-70Hz, to be correlated with excitation to inhibition (E/I) ratio^20^. We applied FOOOF’s linear fitting to LFP data attained using tetrodes in mouse hippocampus, specifically in CA1 and DG sub-regions (see methods, purple and yellow boxes respectively). Due to periodic components in hippocampal ephys signals going as high as 200Hz^21^, we attempted to fit the aperiodic part to a large range of frequencies (4-200Hz). FOOOF’s linear fitting (termed ‘1exp’), gave acceptable fits in smaller frequency ranges (4-100Hz), but got qualitatively and quantitatively worse (see methods – error estimation) when the frequency range was increased to 200Hz (fig 2a - column 1, fig 3a). Further, FOOOF also allows this fitting to have a ‘knee’ frequency, with a flattened fit for lower frequencies, and a linear fit for higher frequencies (termed ‘flat+1exp’, see methods, equation (2)). On using ‘flat+1exp’ fitting, we got more accurate fits for some datasets (fig 2a, column 2), with lower errors (fig 2b, 3a), but some data sets still demanded improvement (fig 2a, row 2).

**Fig 2:**
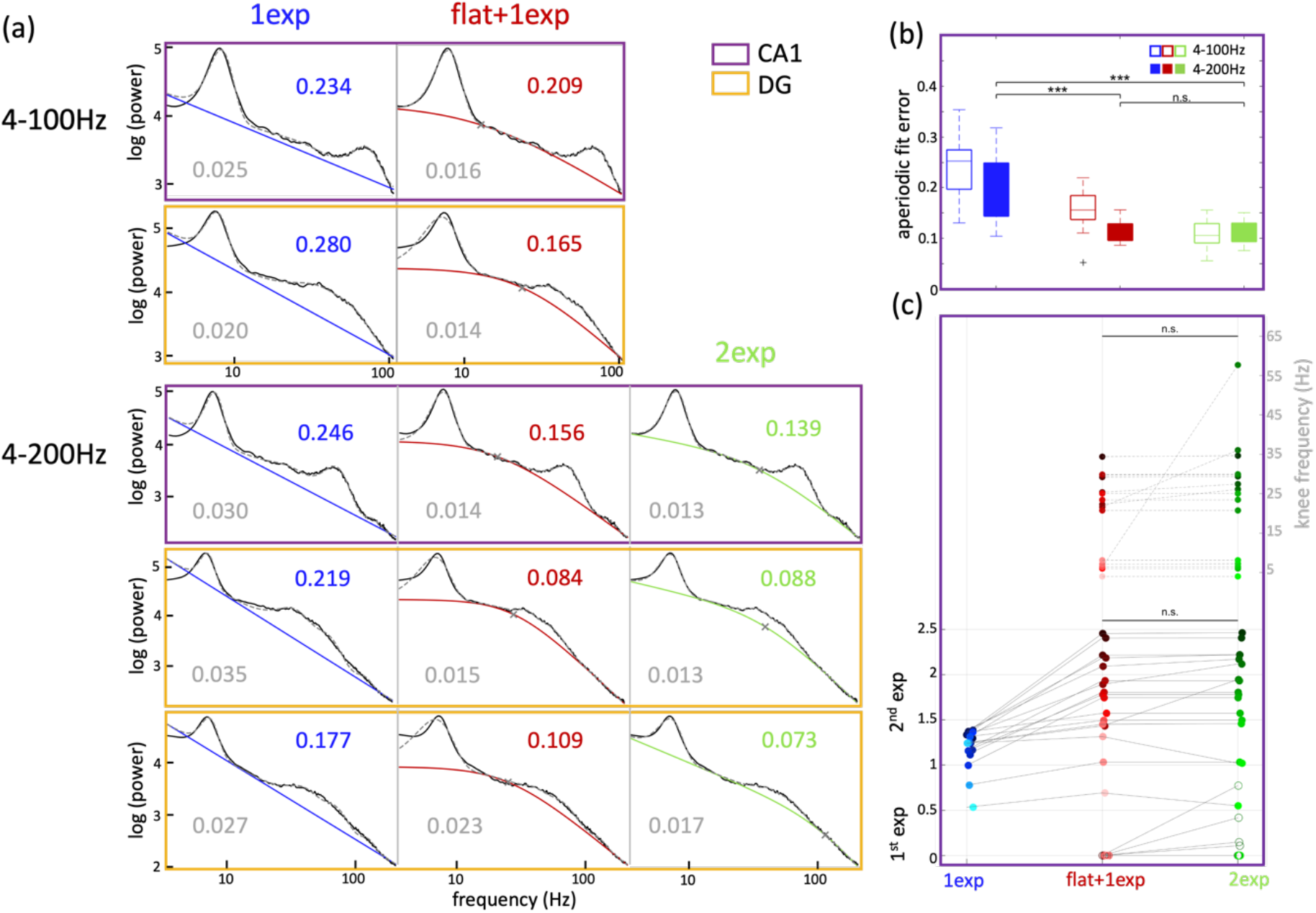
Aperiodic components in hippocampal electrophysiological signals between 4-200Hz. (a) Examples of aperiodic fitting of mouse CA1 (purple) and DG (yellow) electrophysiological signals using ‘1exp’ (blue), ‘flat+1exp’ (red) and ‘2exp’ (green), between 4-100Hz (rows 1-2) or 4-200Hz (rows 3-5). Aperiodic fit errors are shown in color on top right for respective fitting methods, and the corresponding periodic fit errors are show in grey on bottom left. Row 1 and 3 show the same dataset, as do Row 2 and 4. (b) Aperiodic fit errors for CA1 signals (n=17) across ‘1exp’, ‘flat+1exp’ and ‘2exp’, when fit in the ranges of 4-100Hz (lighter colors) and 4-200Hz (darker colors). 4-200Hz paired t-test, p(‘1exp’|’flat+1exp’) = 2.73X10^-5^, p(‘1exp’|’2exp’) = 1.16X10^-5^, p(‘flat+1exp’|’2exp’) = 0.054. (c) Change in the aperiodic components for CA1 signals between the three fitting methods. Paired t-test, 2^nd^ exponent: p(‘flat+1exp’|’2exp’) = 0.48, ‘knee’ frequency: p(‘flat+1exp’|’2exp’) = 0.18. For all figures, ^***^ p<0.0005, ^**^ p<0.005, ^*^ p<0.05, n.s. not significant.

**Fig 3:**
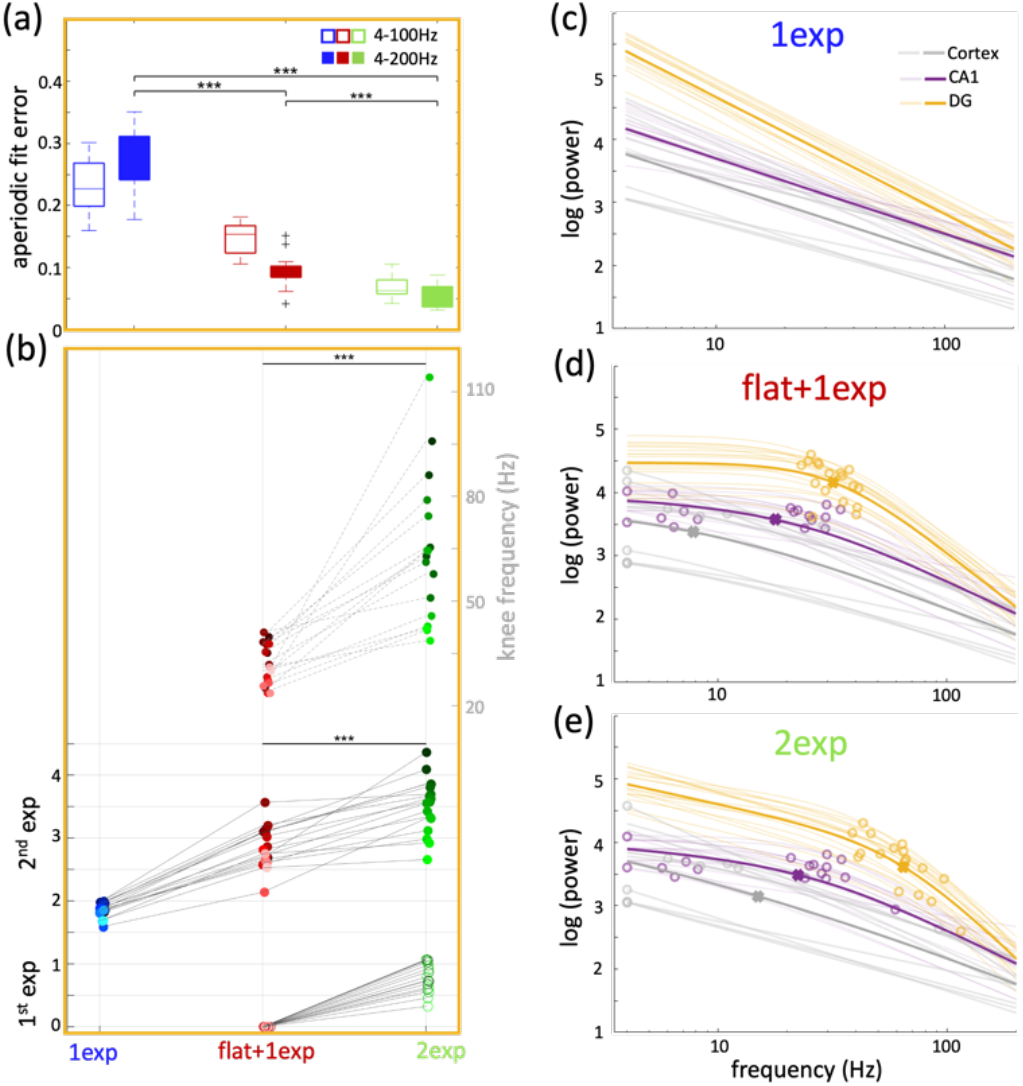
Comparison of aperiodic components between CA1 and DG. (a) Aperiodic fit errors for DG signals across ‘1exp’, ‘flat+1exp’ and ‘2exp’, when fit in the ranges of 4-100Hz (lighter colors) and 4-200Hz (darker colors). 4-200Hz paired t-test, p(‘1exp’|’flat+1exp’) = 6.63X10^-11^, p(‘1exp’|’2exp’) = 1.01X10^-11^, p(‘flat+1exp’|’2exp’) = 7.25X10^-6^. (b) Change in the aperiodic components for DG signals (n=17) between the three fitting methods. paired t-test, 2^nd^ exponent: p(‘flat+1exp’|’2exp’) = 3.76X10^-6^, ‘knee’ frequency: p(‘flat+1exp’|’2exp’) = 1.47X10^-5^. (c) Aperiodic fits using ‘1exp’ of all data from DG (yellow, n=17), CA1 (purple, n=17) and cortex (grey, n=10) with the respective ‘knee’ frequencies as circles. The average of all fits is shown in bold with the ‘knee’ frequency as crosses. Unpaired t-test, exponent: p(DG|CA1) = 4.8X10^-12^, p(CA1|Cortex) = 0.81. (d) Same as (c) but using ‘flat+1exp’. Unpaired t-test, exponent: p(DG|CA1) = 5.71X10^-9^, p(CA1|Cortex) = 0.06, ‘knee’ frequency: p(DG|CA1) = 5.38X10^-5^, p(CA1|Cortex) = 0.02. (e) Same as (c) but using ‘2exp’. Unpaired t-test, 2^nd^ exponent: p(DG|CA1) = 6.33X10^-12^, p(CA1|Cortex) = 0.147, 1^st^ exponent: p(DG|CA1) = 1.86X10^-10^, p(CA1|Cortex) = 0.388, ‘knee’ frequency: p(DG|CA1) = 1.45X10^-7^, p(CA1|Cortex) = 0.374.

As this method was originally optimized for EEG and ECoG signals, both of which are limited in sampling frequency^4^, restraining the maximum fitting frequency to 100Hz would work well. To optimize this method to analyze ephys data and fit the signals to a large range of frequencies, we established new aperiodic fits. Observationally, ephys signals in 4-200Hz were composed of two linearly varying bits with a varying ‘knee’ frequency. We thus established a new fitting paradigm, termed ‘2-exponents’ (‘2exp’, see methods, equation (3)), providing the possibility for the data before the ‘knee’ to have a non-zero exponent as well. Using this fitting function, we got better qualitative fits (fig 2a, column 3), and quantitatively lower aperiodic errors (fig 2b, 3a). Applying ‘2exp’ fits to hippocampal ephys signals only marginally refined the fitting errors for CA1 (fig 2b, supp fig 2a), but it substantially improved the aperiodic fit errors improved by about 50% and periodic fit errors by about 25%, (fig 3a, supp fig 2b) for DG signals, implying differential biological signatures between these regions. We found all aperiodic components from CA1 to be multi-fold lower than in the DG (fig 3e, mean CA1 parameters: 1^st^ exp = 0.09, 2^nd^ exp = 1.76, knee = 22.2Hz; mean DG parameters: 1^st^ exp = 0.79, 2^nd^ exp = 3.52, knee = 64.0Hz). The CA1 parameters were found to not be significantly different than in cortical ephys data (fig 3c,d,e, mean cortex parameters: 1^st^ exp = 0.19, 2^nd^ exp = 1.42, knee = 14.9Hz). Some of these aperiodic parameters follow similar trends when using the ‘1exp’ and ‘flat+1exp’ fits as well (fig 3c,d), but their exact values might be inaccurate due to worse fitting errors.

Biological significance of these aperiodic parameters can be gleaned from previous literature. A recent study modeled the 1/f exponent *n* to also be correlated with the excitation-inhibition (E/I) balance^20^. Using LFP data recorded in rats’ CA1 via shank electrodes, and ECoG data from macaques, they showed that increasing E/I ratio flattened the exponent *n*. As is expected from literature^22,23^, with higher excitation and lower inhibition in CA1, the E/I ratio should be higher in CA1, leading to flatter 1/f slopes in CA1 than in the DG, though which of the two exponents would be more indicative of this ratio is unclear. On the other hand, the ‘knee’ can be thought of as inversely related to different timescales in the data, which reflects the speed of synaptic and transmembrane currents^12,24^. In humans and macaques this is found to increase along the sensorimotor-to-association cortical axis^24^. An increase in this timescale leads to a decrease in the ‘knee’ frequency, leading to an inverse correlation in the ‘knee’ frequency along this axis. Between the hippocampal sub-regions, higher propensity of complex burst spiking in CA1 than in DG should correlate with longer timescales and lower ‘knee’ frequencies in CA1 than DG^25^. The ‘knee’ frequencies found in the mice hippocampal sub-regions translate to timescales of about 7.2ms in CA1 and 2.5ms in DG, aligning with the known bursting characteristics of these regions^26^. Thus, the aperiodic parameters we obtained within the hippocampal sub-regions are coherent with the differences between these sub-regions as reported in the literature.

### Aperiodic fitting until 500Hz

Periodic components in ephys recordings often extend beyond 200Hz, such as sharp-wave ripples in CA1 and CA3^21^. Due to this, some example signals had qualitatively worse fits and higher fitting errors until 200Hz (‘broken examples’, fig 4a, top row). Thus, to obtain complete fits of the periodic components, we further fit our data until 500Hz (see methods). Applying ‘2-exp’ aperiodic fitting to hippocampal mouse data until 500Hz, gave much poorer fits with higher errors (fig 4a,c,d). To improve this, we further developed two more aperiodic fits to cover the whole range of frequencies from 4-500Hz. To match the observational aperiodic nature of the data extending into these higher frequencies, we added a third linear piece to the above curves, adding a second ‘knee’ and potentially a third exponent. For the first of these fits, we fixed the third exponent to be 0, i.e. have a flat end to the spectrum, thus naming this fit as ‘2exp+flat’. We also used another fit, with the third exponent also be a free fitting parameter, thus naming this ‘3exp’ (see methods, equations (4,5)).

**Fig 4:**
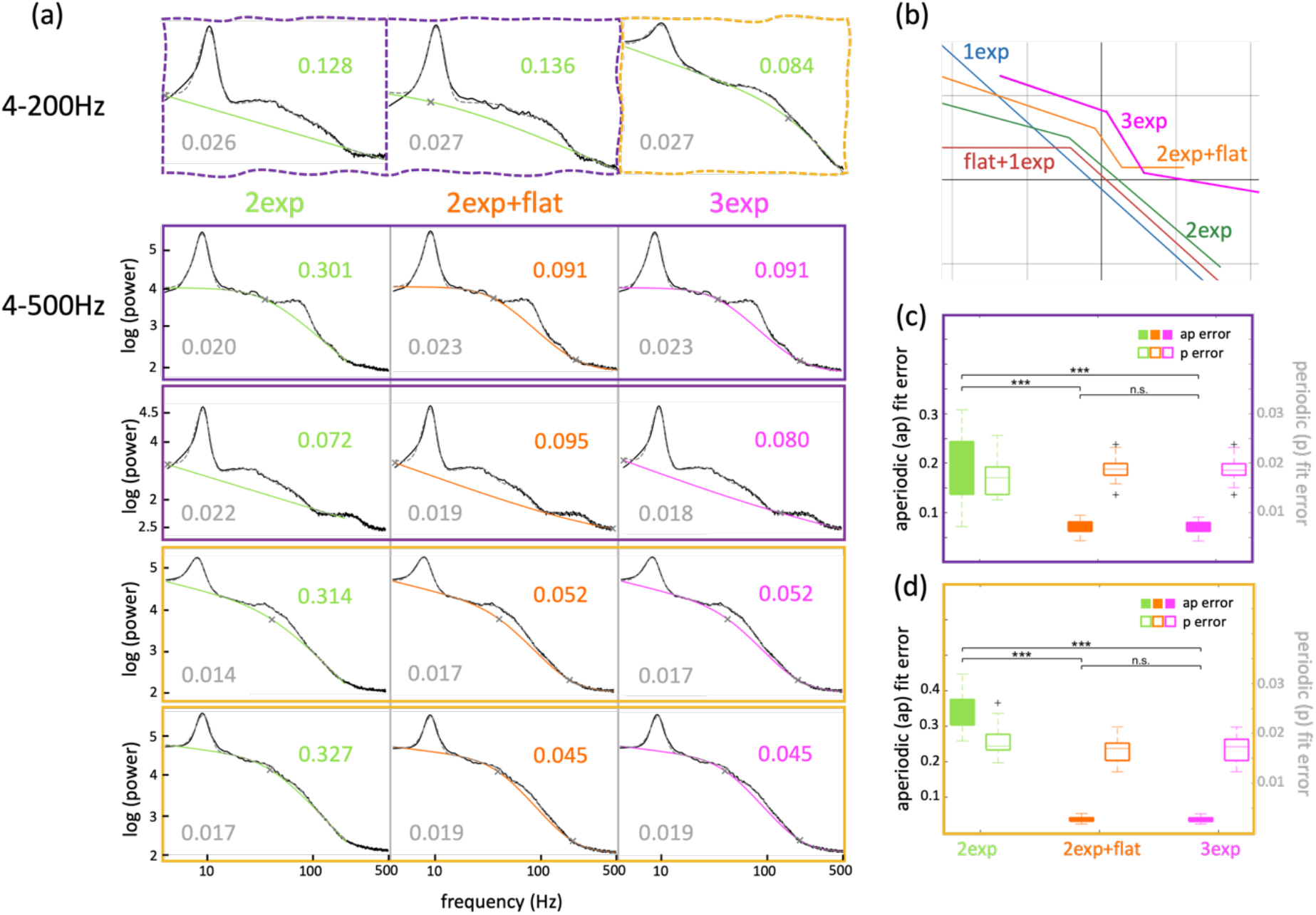
Aperiodic components in hippocampal electrophysiological signals between 4-500Hz. (a) Row 1 shows ‘broken examples’ of aperiodic fitting of mouse CA1 (purple) and DG (yellow) electrophysiological signals using ‘2exp’ between 4-200Hz. Rows 2-5 show similar examples using ‘2exp’ (green), ‘2exp+flat’ (orange) and ‘3exp’ (magenta), between 4-500Hz. Row 1, column 1 and Row 3 show the same dataset, Row 4 and Row 2 and 4 from fig 2 show the same dataset. (b) The general form of all 5 fitting functions. (c) Aperiodic (darker colors, left y-axis) and periodic (lighter colors, right y-axis) fit errors for CA1 signals across ‘2exp’, ‘2exp+flat’ and ‘3exp’, when fit in 4-500Hz. Paired t-test, Aperiodic errors: p(‘2exp’|’2exp+flat’) = 1.95X10^-6^, p(‘2exp’|’3exp’) = 1.25X10^-6^, p(‘2exp+flat’|’3exp’) = 0.32. (d) Same as (c) but for DG signals. Paired t-test, Aperiodic errors: p(‘2exp’|’2exp+flat’) = 1.43X10^-13^, p(‘2exp’|’3exp’) = 1.76X10^-13^, p(‘2exp+flat’|’3exp’) = 0.07.

Fitting until higher frequencies helped with better estimates of the periodic components, such as higher power in specific oscillations or detectability of very low power oscillations (fig 4a). Examples that couldn’t be fit well with previous models (‘broken examples’, fig 4a, top row, column 1), fit much better at higher frequencies with the two new models (fig 4a, second row), and errors got better with both these models (fig 4c,d). The general form of all the fitting functions is shown in fig 4b. Though for majority of the cases, both ‘2exp+flat’ and ‘3exp’ functions behave similarly, some cases preferentially fit better with one or the other (fig 4a, second row), providing flexibility in fitting a wide variety of signals without prior assumptions.

To compare the efficacy of the fitting models, we compared the Bayesian information criterion (bic) scores (supp fig 2c, see methods, equation (6)) for the first 3 models between 4-200Hz, and for the last 2 models between 4-500Hz. The change in bic scores clearly show that the more complex models fit the data better, with the increasing complexity being particularly critical for the analyses of DG signals.

## DISCUSSION

With crucial modifications to the open-source toolbox FOOOF, we decreased the estimation errors for periodic components while also assessing the aperiodic components of mouse hippocampal electrophysiological signals, not only for a narrow range of frequencies, but also within a wide range from 4-500 Hz.

For the assessment of the periodic components, using the conventional methods that do not estimate the aperiodic fits of the neural data, such as band-pass filtering or Chronux toolbox, the estimates of the cf of the brain oscillations will always be underestimated (fig 1b). This underestimation would further increase for oscillations with lower frequencies and at higher aperiodic exponents. Correctly estimating these frequencies is essential to compare conditions that cause only small changes in the brain rhythms. Even though the improvement in errors with our iterative FOOOF procedure was seemingly small (3-5%), it is comparable to the changes in theta cf associated with behavioral parameters, such as speed and acceleration^5^. As the frequencies of these rhythms are also thought to be correlated with the speed of inter-regional synchrony^27^, small changes in these might have important biological significance. Our methods would thus be crucial in accurately correlating changes in brain rhythms with animal’s cognitive states or behavioral contingencies.

Our improved toolbox was critical to compare the aperiodic parameters between different hippocampal sub-regions. LFP and ECoG signals have been reported to have relatively constant spectral power at low frequencies (1-10Hz)^20^. For tetrode-acquired hippocampal ephys signals, this was found to be only partially true. At these low frequencies, the exponents were found to small (<1), but non-zero for a few CA1 signals (4/17), while they were always non-zero for all DG signals (1^st^ exp comparisons fig 2c, 3b). For most of the signals from CA1, these aperiodic parameters did not change at all or by only a little between the ‘flat+1exp’ and ‘2exp’ methods (fig 2c). Conversely, they changed significantly for the DG signals (fig 3b). This shows that while the aperiodic fitting as done within the original FOOOF toolbox might work for EEG and hippocampus CA1 ephys data, our modifications are essential for DG ephys data. This is also evident in the change in bic scores (supp fig 2c). It is important to note that with ‘1exp’ fitting, the exponents being lower for CA1 as compared with DG can be trivially due to higher starting power in the DG, as the power that all signals fall down to would depend on the acquisition system. This might also play a role in comparing the 2^nd^ exponent with the ‘flat+1exp’ and ‘2exp’ functions, but the comparisons between the 1^st^ exponent and the ‘knee’ frequency should stay unbiased by that.

Recent reports suggest that the E/I ratio might not be uniform throughout the DG, with its upper blade having higher inhibition than its lower blade^28,29^. Our analyses methods as outlined in this work can be valuable in estimating differences between the E/I ratios of the two DG blades. Further, our methods will be crucial to assess the changes in various periodic and aperiodic parameters, such as center frequency, exponents and ‘knee’ frequencies, to various behavioral and cognitive states, such as during hippocampal memory tasks, sleep replay, aging, performance in virtual reality^30^, etc.

## Supplementary figures

**Supp Fig 1:**
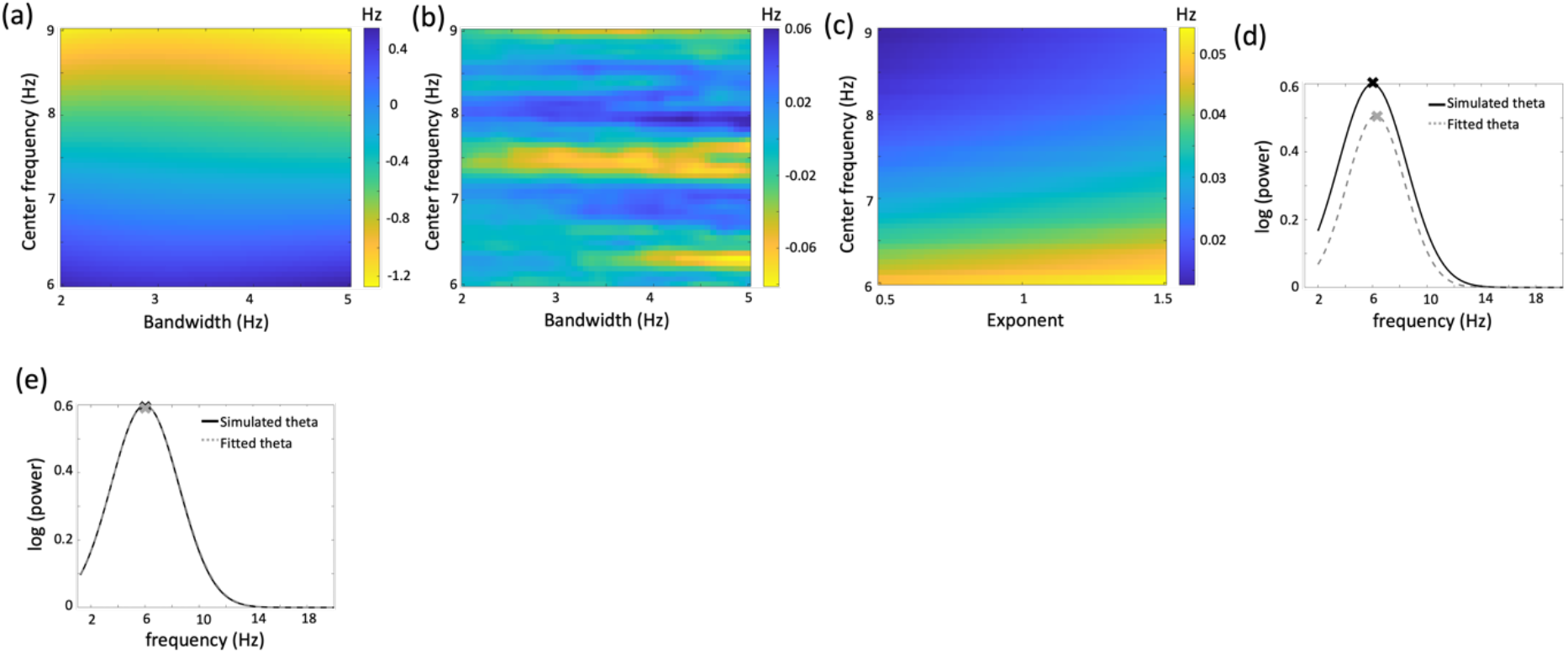
Better estimates of the periodic components in simulated electrophysiological signals. (a) Errors in cf estimates of the simulated theta peak with pink-noise exponent of 1.2, using band-pass filtering. (b) Errors in cf estimates of the simulated theta peak with pink-noise exponent of 0, using Chronux. This was smoothed using a Gaussian filter with std of 1. (c) Average errors in cf estimates using FOOOF, across 20 randomizations, of the simulated theta peak (with bw of 3.5Hz), with varying exponent and cf. (d) Flattened simulated and assessed theta Gaussian peak (cf = 6Hz, bw = 5Hz, exponent = 1.2) using FOOOF, without iterations. (e) Same as (d) but after 20 iterations.

**Supp Fig 2:**
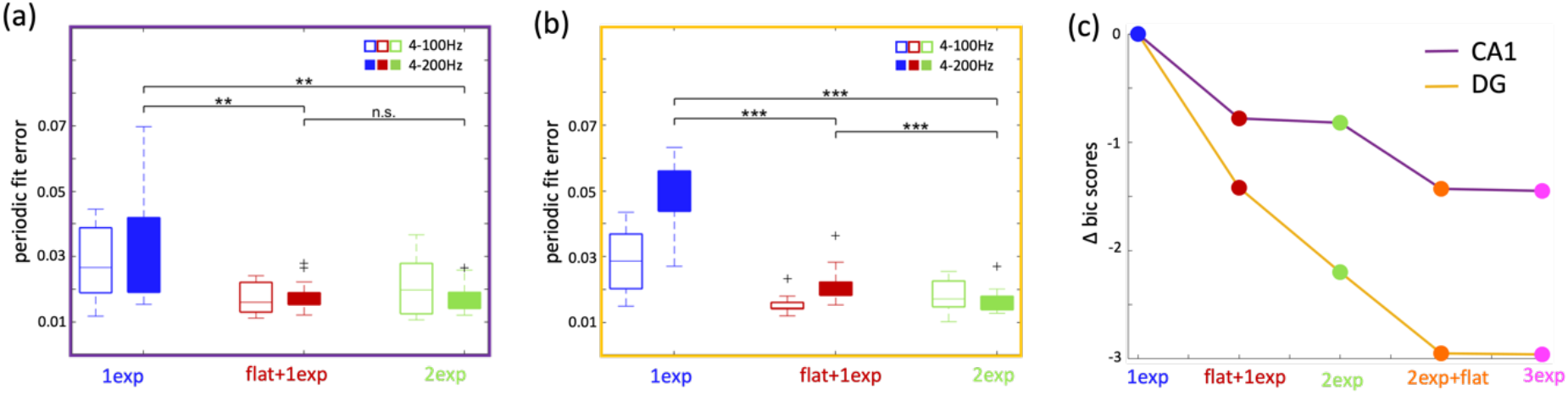
Periodic fit errors. (a) Periodic fit errors for CA1 signals across ‘1exp’, ‘flat+1exp’ and ‘2exp’, when fit in the ranges of 4-100Hz (lighter colors) and 4-200Hz (darker colors). 4-200Hz paired t-test, p(‘1exp’|’flat+1exp’) = 0.001, p(‘1exp’|’2exp’) = 9.4X10^-4^, p(‘flat+1exp’|’2exp’) = 0.07. (b) Same as (a) but for DG signals. 4-200Hz paired t-test, p(‘1exp’|’flat+1exp’) = 4.17X10^-8^, p(‘1exp’|’2exp’) = 1.32X10^-9^, p(‘flat+1exp’|’2exp’) = 5.69X10^-5^. (c) The change in bic scores across the 5 aperiodic fitting models, averaged over all CA1 and DG signals.

## Methods

### Subjects

Three adult male C57BL/6JRj mice (Janvier Labs) (approximately 3-4 months old at the time of surgical implantation) were individually housed on a 12-hour dark/light (reverse) cycle. Animals were given ad-lib food and water. All experimental procedures were approved by local authorities and compiled with all relevant ethical regulations (TVV 49/2022).

### Animal Handling and Surgery

The mice were handled for around 15 days to get them used to the room, experimenters, and being gently restrained within and between the hands of the experimenters. The animals that stayed calm, yet curious, during these conditions were then considered for implantation. The mice were implanted with 2-3g ShuttleDrives from OpenEphys (Voigts, Harnett, 2019), with 12-14 individually adjustable tetrodes (13um nichrome wires) positioned unilaterally on the right hemisphere over dorsal CA1 (-2.0mm A.P., 1.7mm M.L. relative to bregma). Surgery was performed under isoflurane anesthesia and heart rate, breathing rate, and body temperature were continuously monitored.

Analgesia was achieved by using Metamizol (200mg/kg, sc), Buprenovet (0.1mg/kg, sc) and Xylocaine Gel (2%, topical). One 1.6mm-diameter craniotomy was drilled using a hand-held drill. Dura mater was removed and the ShuttleDrive was lowered until the cannula is level with the skull surface. The implant was anchored to the skull with 6 skull screws and dental cement. One parietal and one occipital skull screw was used as ground for recording. Mice were administered 150mg Metamizol in 100ml drinking water (200mg/kg) for 7 days during recovery.

### Environment and Animal Behavior

All experiments were conducted inside a 1.5mX1.5mX1.5m EMF-shielded enclosure, with only red lights to maintain the reverse light cycle. The animals ran in an open-field environment (OFE), a 50cmX50cmX50cm box, while they foraged for randomly scattered chocolate sprinkles. The base of the OFE box was covered with a white uniform smooth mat, which was wiped clean at the end of each session with ethanol to minimize scents between days. The animal’s behavior is captured with a 55Hz color Chameleon3 FLIR camera, and synced with the neural data using Bonsai (Lopes, Kampff, 2015).

### Electrophysiology and Neural Signal Acquisition

The tetrodes were lowered gradually after surgery into the hippocampus. Positioning of the electrodes in CA1 and DG was confirmed through the presence of sharp-wave ripples and dentate spikes during recordings, and through histology after experiments. Neural signals were acquired using Intan recording systems (C3100, sampling rate 30kHz) and visualized on Open Ephys GUI, using OpenEphys low-profile SPI headstage 64ch.

### Simulated data

All the parameters for simulating the neural signals were chosen to emulate our tetrode ephys hippocampal signals. We started with Matlab ‘randn’ function, added 1/f^n^ with n of 1.2 to it for the whole range of 1-1000Hz, and put in an offset such that the power at 1Hz is 60dB. The exponent of 1.2 chosen here was slightly higher than the maximum for real DG data (∼1.1, median∼0.7, fig 3b) to heighten the fallacies in FOOOF, especially when the periodic components are at lower frequencies. Theta peak was put in as a Gaussian peak with a center frequency (cf) of 6-9Hz, maximum power of about 6dB above the cf, and a standard deviation of 1-2.5Hz (2-5Hz bandwidth (bw)). Similarly, to simulate peaks in the sgamma band, we put in a Gaussian peak with cf of 30-40Hz, maximum power of about 2dB above cf, and bw of 5-15Hz bw. We also put in a 2.5dB, 0.5Hz bw noise peak centered at 50Hz, and another one centered at 150Hz with the same parameters.

### Preprocessing of neural data

The ephys signals were downsampled to 1kHz to be used as LFP traces. The segments when the animal was moving below a speed of 5cm/s for at least 450ms were replaced with zeros, to avoid any confounds in the periodic or aperiodic components with sleeping or quiescent states. For each electrode, the power spectral density (psd) was calculated, using the Welch’s method with a Hanning window size of 1.2s (3s for fig 1), nfft of 4000 and 50% overlap. The electrode with the highest total power [∑_1<*F*<495_ log_10_ p*sd*(*F*)] was chosen from each tetrode. Electronic noise peak at 50Hz was removed from the psd by applying the *‘*1exp*’* FOOOF fit in a small range (43-57Hz) and subtracting the fitted Gaussian. The same procedure was applied to eliminate any harmonics of theta and 50Hz noise.

### Equations for the aperiodic fits

Simplified versions of the equations are shown here for clarity, the exact forms can be found in the available code

For 4-100Hz and 4-200Hz, log(power) *L* for each spectrum could be fit to

<u>1exp:</u>

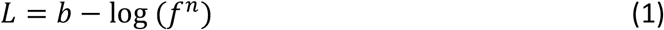

where *n* is the exponent across frequencies *f*

<u>flat+1exp:</u>

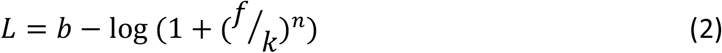

where *k* is the knee frequency in Hz, and *n* is the exponent across frequencies *f*

<u>2exp:</u>

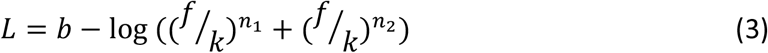

where *k* is the knee frequency in Hz, and *n*_*1*_ *and n*_*2*_ are the exponents across frequencies *f*

For direct comparison of the ‘2exp’ fitting with ‘flat+1exp’ fitting, we put the following conditions on the aperiodic components after the fittings: if the absolute difference between the two exponents is less than 0.01, or if the reported ‘knee’ frequency is greater than 195Hz, we change the 1^st^ exponent to be 0, 2^nd^ exponent to be the as-obtained value and the knee frequency to 4Hz.

For 4-500Hz fitting: We restricted the fitting to 495Hz to eliminate roll-off artifacts

<u>2exp+flat:</u>

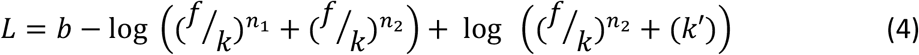

where *n*_*1*_ *and n*_*2*_ are the exponents across frequencies *f* with n_2_>n_1_, k is the 1^st^ knee, (k’/n_2_ + k) is the 2^nd^ knee frequency, both in Hz.

<u>3exp:</u>

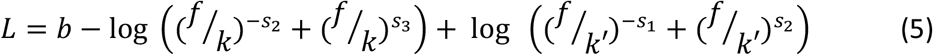

where n_1_ = s_1_-s_2_, n_2_ = s_1_+s_3_ and n_3_ = s_3_-s_2_ are the three exponents, with s_1_, s_2_, s_3_>0, and k and k’ are the 1^st^ and 2^nd^ knee frequencies in Hz, respectively.

To avoid over-fitting until 500Hz, we implemented step-wise curve-fitting, i.e. fit the data until 200Hz with the ‘2exp’ function, fixed the ‘1^st^ exponent’, ‘1^st^ knee frequency’ and starting power at 4Hz, which might be biologically more relevant, and then re-fit the data. This allows the 2^nd^ and 3^rd^ exponent, and the 2^nd^ knee, to be fit freely. This step-wise fitting can prevent mathematical operations from messing up biologically relevant parameters.

### Detection of periodic components

We started both aperiodic and periodic assessment at 4Hz to avoid delta rhythm, which can be inconsistent in the hippocampus. As done in the original FOOOF toolbox^13^, we processed each psd by subtracting the most suitable aperiodic fit from it resulting in a flattened spectrum. Thereafter, instead of using FOOOF’s multi-Gaussian fitting method, we divided the flattened spectrum into hippocampus-relevant oscillatory bands and ran the peak detection process only once per band. This was done to avoid an individual brain rhythm being represented as a combination of multiple Gaussians, which posed a challenge for further analyses and interpretation. This additionally allowed us to define individual thresholds and restrictions for each oscillatory band. The following parameters were used in this process:

| Detection range (Hz) | Cf-bounds (Hz) | Peak-width limits (Hz) | Peak thresholds | Min. peak-height (log10(psd)) |
| --- | --- | --- | --- | --- |
| 4-12 (theta) | 5, 9.5 | 2, 5 | 1 | 0.1 |
| 19-48 (sgamma) | 25, 40 | 5, 25 | 1 | 0.05 |
| 51-90 (fgamma) | 52, 80 | 3, 35 | 1 | 0.08 |
| 100-200 (ripples) | 125, 160 | 10, 30 | 1 | 0.05 |
| 205-495 | 220, 380 | 50, 150 | 1 | 0.05 |

This modification is different from the ‘Band-by-Band’ approach introduced with FOOOF 2.0, namely specparam, which relies on removing unwanted Gaussian peaks in post-analysis.

### Errors estimation

To quantify the goodness of our fittings, we used two kinds of error estimates:

- Aperiodic (ap) fit error: These were calculated to quantify the fittings of the aperiodic components. To do this, we took the mean absolute difference of our data and the aperiodic fits, across all frequency points.
- Periodic (p) fit error: These were calculated to quantify the fittings of the periodic components. To do this, we took the mean absolute difference of our data with the periodic fits obtained with FOOOF, across all frequency points.

As expected, the periodic fit errors are only marginally affected by the fitting modes, as flattening out the spectrum using the aperiodic components already makes the periodic fitting quite good.

### Bayesian information criterion (bic) calculations

bic scores were calculated for all the fitting models using the following equations

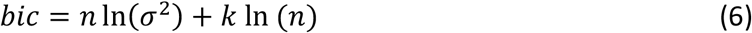

where n is the number of frequency bins, k is the number of free parameters and *σ*^2^ is the mean of the mean square of the ap error for each model.

## REFERENCES

1. Buzsáki, G., Costas A. Anastassiou, and Christof Koch (2012). The origin of extracellular fields and currents—EEG, ECoG, LFP and spikes. Nat Rev Neurosci 13, 407–420. 10.1038/nrn3241.

2. Buzsáki, G., and Draguhn, A. (2004). Neuronal Oscillations in Cortical Networks. Science (1979) 304, 1926–1929. 10.1126/science.1099745.

3. Donoghue, T., Schaworonkow, N., and Voytek, B. (2022). Methodological considerations for studying neural oscillations. European Journal of Neuroscience 55, 3502–3527. 10.1111/ejn.15361.

4. Gerster, M., Waterstraat, G., Litvak, V., Lehnertz, K., Schnitzler, A., Florin, E., Curio, G., and Nikulin, V. (2022). Separating Neural Oscillations from Aperiodic 1/f Activity: Challenges and Recommendations. Neuroinformatics 20, 991–1012. 10.1007/s12021-022-09581-8.

5. Kennedy, J.P., Zhou, Y., Qin, Y., Lovett, S.D., Sheremet, A., Burke, S.N., and Maurer, A.P. (2022). A Direct Comparison of Theta Power and Frequency to Speed and Acceleration. The Journal of Neuroscience 42, 4326–4341. 10.1523/JNEUROSCI.0987-21.2022.

6. Leung, S.L. (1998). Generation of Theta and Gamma Rhythms in the Hippocampus. Neurosci Biobehav Rev 22, 275–290. 10.1016/S0149-7634(97)00014-6.

7. Nunez, P.L. (1981). A Study of Origins of the Time Dependencies of Scalp EEG: I -Theoretical Basis. IEEE Trans Biomed Eng BME-28, 271–280. 10.1109/TBME.1981.324700.

8. Buzsáki, G., Logothetis, N., and Singer, W. (2013). Scaling Brain Size, Keeping Timing: Evolutionary Preservation of Brain Rhythms. Neuron 80, 751–764. 10.1016/j.neuron.2013.10.002.

9. Bokil, H., Andrews, P., Kulkarni, J.E., Mehta, S., and Mitra, P.P. (2010). Chronux: A platform for analyzing neural signals. J Neurosci Methods 192, 146–151. 10.1016/j.jneumeth.2010.06.020.

10. Donoghue, T., Dominguez, J., and Voytek, B. (2020). Electrophysiological Frequency Band Ratio Measures Conflate Periodic and Aperiodic Neural Activity. eNeuro 7, ENEURO.0192-20.2020. 10.1523/ENEURO.0192-20.2020.

11. Safaryan, K., and Mehta, M.R. (2021). Enhanced hippocampal theta rhythmicity and emergence of eta oscillation in virtual reality. Nat Neurosci 24, 1065–1070. 10.1038/s41593-021-00871-z.

12. Donoghue, T., Hammonds, R., Lybrand, E., Washcke, L., Gao, R., and Voytek, B. (2024). Evaluating and Comparing Measures of Aperiodic Neural Activity. BioRXiv. 10.1101/2024.09.15.613114.

13. Donoghue, T., Haller, M., Peterson, E.J., Varma, P., Sebastian, P., Gao, R., Noto, T., Lara, A.H., Wallis, J.D., Knight, R.T., et al. (2020). Parameterizing neural power spectra into periodic and aperiodic components. Nat Neurosci 23, 1655–1665. 10.1038/s41593-020-00744-x.

14. Kala, A., Leemburg, S., and Jezek, K. (2023). Sepsis-Induced Changes in Spectral Segregation and Kinetics of Hippocampal Oscillatory States in Rats. eNeuro 10, ENEURO.0002-23.2023. 10.1523/ENEURO.0002-23.2023.

15. Leemburg, S., Kala, A., Nataraj, A., Karkusova, P., Baindur, S., Suresh, A., Blahna, K., and Jezek, K. (2024). LPS-induced sepsis disrupts brain activity in a region- and vigilance-state specific manner. BioRXiv. 10.1101/2024.12.25.630319.

16. Ravassard, P., Kees, A., Willers, B., Ho, D., Aharoni, D., Cushman, J., Aghajan, Z.M., and Mehta, M.R. (2013). Multisensory control of hippocampal spatiotemporal selectivity. Science (1979) 340, 1342–1346.

17. Belluscio, M.A., Mizuseki, K., Schmidt, R., Kempter, R., and Buzsáki, G. (2012). Cross-Frequency Phase–Phase Coupling between Theta and Gamma Oscillations in the Hippocampus. The Journal of Neuroscience 32, 423–435. 10.1523/JNEUROSCI.4122-11.2012.

18. Strüber, M., Sauer, J.-F., and Bartos, M. (2022). Parvalbumin expressing interneurons control spike-phase coupling of hippocampal cells to theta oscillations. Sci Rep 12, 1362. 10.1038/s41598-022-05004-5.

19. Boncompte, G., Medel, V., Irani, M., Lachaux, J.P., and Ossandon, T. (2024). Aperiodic exponent of brain field potentials is dependent on the frequency range it is estimated. BioRXiv. 10.1101/2024.12.17.628966.

20. Gao, R., Peterson, E.J., and Voytek, B. (2017). Inferring synaptic excitation/inhibition balance from field potentials. Neuroimage 158, 70–78. 10.1016/j.neuroimage.2017.06.078.

21. Buzsáki, G. (2015). Hippocampal sharp wave-ripple: A cognitive biomarker for episodic memory and planning. Hippocampus 25, 1073–1188. 10.1002/hipo.22488.

22. Arima-Yoshida, F., Watabe, A.M., and Manabe, T. (2011). The mechanisms of the strong inhibitory modulation of long-term potentiation in the rat dentate gyrus. European Journal of Neuroscience 33, 1637–1646. 10.1111/j.1460-9568.2011.07657.x.

23. Alkadhi, K.A. (2019). Cellular and Molecular Differences Between Area CA1 and the Dentate Gyrus of the Hippocampus. Mol Neurobiol 56, 6566–6580. 10.1007/s12035-019-1541-2.

24. Gao, R., van den Brink, R.L., Pfeffer, T., and Voytek, B. (2020). Neuronal timescales are functionally dynamic and shaped by cortical microarchitecture. Elife 9. 10.7554/eLife.61277.

25. Galashin, A.S., Konakov, M. V., and Dynnik, V. V. (2024). Comparison of Spontaneous and Evoked Activity of CA1 Pyramidal Cells and Dentate Gyrus Granule Cells of the Hippocampus at an Increased Extracellular Potassium Concentration. Biochem (Mosc) Suppl Ser A Membr Cell Biol 18, 339–347.

26. Harris, K.D., Hirase, H., Leinekugel, X., Henze, D.A., and Buzsáki, G. (2001). Temporal Interaction between Single Spikes and Complex Spike Bursts in Hippocampal Pyramidal Cells. Neuron 32, 141–149. 10.1016/S0896-6273(01)00447-0.

27. Ahmed, O.J., and Mehta, M.R. (2012). Running Speed Alters the Frequency of Hippocampal Gamma Oscillations. The Journal of Neuroscience 32, 7373–7383. 10.1523/JNEUROSCI.5110-11.2012.

28. Berdugo-Vega, G., Dhingra, S., and Calegari, F. (2023). Sharpening the blades of the dentate gyrus: how adult-born neurons differentially modulate diverse aspects of hippocampal learning and memory. EMBO J 42. 10.15252/EMBJ.2023113524.

29. Luna, V.M., Anacker, C., Burghardt, N.S., Khandaker, H., Andreu, V., Millette, A., Leary, P., Ravenelle, R., Jimenez, J.C., Mastrodonato, A., et al. (2019). Adult-born hippocampal neurons bidirectionally modulate entorhinal inputs into the dentate gyrus. Science (1979) 364, 578–583. 10.1126/science.aat8789.

30. Purandare, C.S., Dhingra, S., Rios, R., Vuong, C., To, T.T., Hachisuka, A., Choudhary, K., and Mehta, M.R. (2022). Moving bar of light evokes vectorial spatial selectivity in the immobile rat hippocampus. Nature 602, 461–467.

